# FiberNeat: Unsupervised White Matter Tract Filtering

**DOI:** 10.1101/2021.10.26.465991

**Authors:** Bramsh Qamar Chandio, Tamoghna Chattopadhyay, Conor Owens-Walton, Julio E. Villalon Reina, Leila Nabulsi, Sophia I. Thomopoulos, Eleftherios Garyfallidis, Paul M. Thompson

## Abstract

Whole-brain tractograms generated from diffusion MRI digitally represent the white matter structure of the brain and are composed of millions of streamlines. Such tractograms can have false positive and anatomically implausible streamlines. To obtain anatomically relevant streamlines and tracts, supervised and unsupervised methods can be used for tractogram clustering and tract extraction. Here we propose FiberNeat, an unsupervised white matter tract filtering method. FiberNeat takes an input set of streamlines that could either be unlabeled clusters or labeled tracts. Individual clusters/tracts are projected into a latent space using nonlinear dimensionality reduction techniques, t-SNE and UMAP, to find spurious and outlier streamlines. In addition, outlier streamline clusters are detected using DBSCAN and then removed from the data in streamline space. We performed quantitative comparisons with expertly delineated tracts. We ran FiberNeat on 131 participants’ data from the ADNI3 dataset. We show that applying FiberNeat as a filtering step after bundle segmentation improves the quality of extracted tracts and helps improve tractometry.

## I. INTRODUCTION

The structural architecture of the brain can be computationally reconstructed from a diffusion magnetic resonance imaging (MRI) [1] dataset using tractography algorithms [2]. Tractography algorithms exploit the direction and paths of water diffusion in neural connections of the brain to generate digital neural pathways, otherwise called streamlines. Streamlines are thus used as a computational approximation of the brain’s white matter fibers. Tractography algorithms often generate streamlines that are false positives or anatomically implausible, such as streamlines that loop, that have sharp curves and angles, that terminate prematurely in white matter, or that connect anatomically implausible regions of the brain [3], [4].

In the past two decades, researchers have used both supervised and unsupervised white matter tract segmentation methods to reduce the number of false positive streamlines in the data. The unsupervised category focuses on clustering methods [5], [6] that divide whole-brain tractograms into clusters of streamlines that are spatially similar in shape and size. Resultant clusters often suffer from spurious streamlines or poor alignment with neuroanatomical definitions of the tracts. Furthermore, clustering methods do not provide anatomically relevant labels to clusters and can have sub-clusters within one cluster. The supervised category consists of white matter tract segmentation methods that are trained with pre-labeled datasets. Automatic tract segmentation methods include ROI-based [7], atlas-based [8], [9], and deep learning-based methods [10]. Although such supervised methods result in labeled streamlines that match their anatomical tract definitions, they can still produce spurious streamlines due to biases stemming from limitations of the prior anatomical reference, subject variability, and tractography reconstruction issues. Moreover, different tract segmentation methods may rely on different definitions of the same tracts [11].

In this paper, we propose FiberNeat, a method which uses dimensionality reduction techniques t-SNE (t-distributed stochastic neighbor embedding) [12] and UMAP (uniform manifold approximation and projection) [13] to find and remove outlier streamlines in latent space ^1^. The input to FiberNeat is a set of streamlines that can either be anatomically unlabeled clusters of streamlines or anatomically labeled tracts. It populates an *N* × *N* square distance matrix by calculating pair-wise distances among all *N* streamlines in the cluster/tract using the streamline based minimum direct-flip distance (MDF) metric [6]. We chose MDF distance metric as a solution to the inconsistent streamline orientation problem. MDF is one of the fastest streamline distance metrics [29] which helps in reducing overall computational time of FiberNeat method. The distance matrix is fed to nonlinear dimensionality reduction methods, i.e., t-SNE or UMAP, to project data into 2D space. In 2D space, spatially close streamlines are placed together and spurious streamlines are placed far from others. Hence, it becomes easier to visually and algorithmically filter out outlier clusters in the latent space. FiberNeat uses the density-based clustering method DBSCAN [14] to computationally label clusters in 2D space. It only keeps the streamlines of the largest clusters and removes small outlier clusters of streamlines. We use labels of small clusters given by DBSCAN in 2D space to remove corresponding clusters of streamlines in streamline-space. FiberNeat is an unsupervised data-driven algorithm that does not require any anatomical reference atlas or labeled training data.

## II. Methods

Input to FiberNeat can be individual clusters from a whole-brain tractogram or extracted white matter tracts, where cluster/tract *C* is a set of *N* streamlines. *C* = {*S*_1_, *S*_2_, …, *S*_*n*_}, *S*_*i*_ ∈*C, S*_*i*_ = {*s*_1_, *s*_2_, …, *s*_*n*_}, where *s*_*i*_ is a 3D vector point. The number of points per streamline may vary.

The FiberNeat method consists of the following steps:

1. Set all streamlines to have *k* number of points
2. Populate *N* × *N* distance matrix *D* by calculating pairwise MDF distances among all streamlines in the set *C*.
3. Project streamlines into 2D space using the precomputed streamline distance matrix *D*.
  - Use either t-SNE or UMAP for the dimensionality reduction.
4. Cluster the streamlines in the 2D latent space using DBSCAN. Smaller clusters of 2D points are considered outliers. Streamlines belonging to the largest cluster in 2D space are kept in streamline space; streamlines belonging to the small clusters are removed.

Fig.1 illustrates steps of the FiberNeat method. A.a is an input set of streamlines that could either be an unlabeled cluster or a labeled white matter bundle. We project individual clusters/tracts into lower dimensional space using t-SNE (A, B). We take an individual cluster of streamlines (A.a, A.b) and calculate pair-wise streamline distances within that cluster (A.c) using the streamline-based MDF distance metric [6]. The MDF distance metric takes into account that streamlines traversing the brain in the same direction can be saved with opposite orientation. This step calculates a direct distance between two streamlines with their default orientation and a distance between a streamline and a streamline with a flipped orientation and selects the minimum of two. We provide t-SNE with this pre-calculated distance matrix as it embeds relevant information on similarities and differences between pairs of streamlines. As both t-SNE and UMAP are manifold learning approaches for non-linear dimensionality reduction, t-SNE could also be replaced by UMAP in this case. While the former captures and preserves local structure, the latter aims to preserve both local and global structure in the data. Streamlines are projected into 2D space by t-SNE (B.a) and the results are then clustered using the density-based clustering method, DBSCAN (B.b). This helps to visually and algorithmically locate outlier streamlines, as those tend to be placed and clustered together (B.b). Class 0 and 2 show outlier streamlines and are filtered out from the initial cluster (A.b) in streamline space (B.c). The entire process is completely unsupervised with no external information provided about anatomy. Visually, (B.c) agrees well with the expected trajectory of the arcuate fasciculus bundle in the left hemisphere of the brain.

**Fig. 1.**
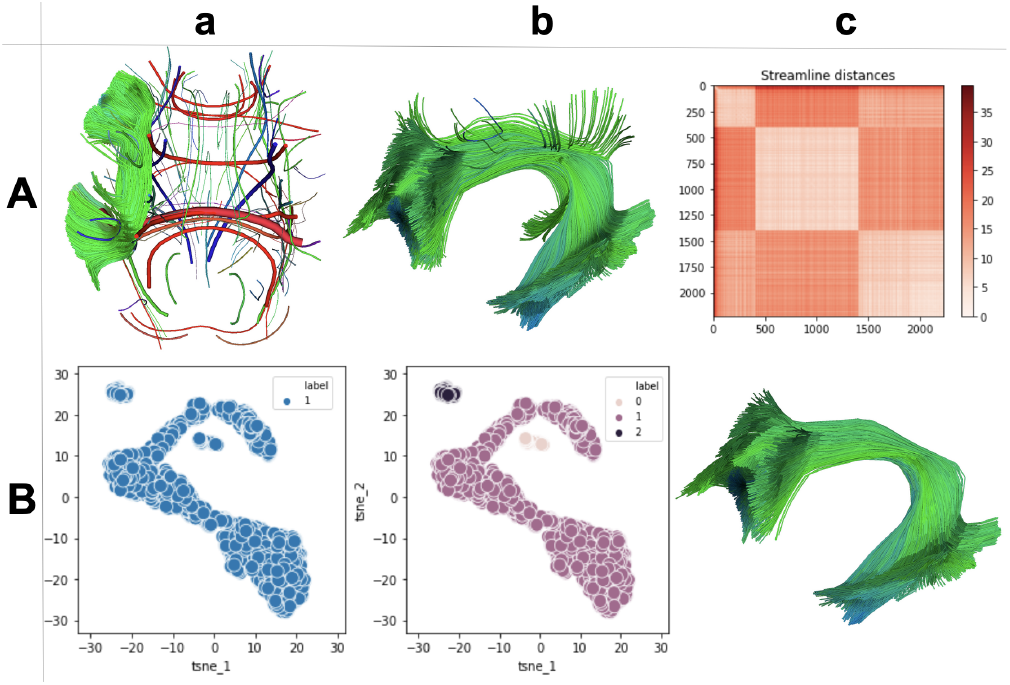
Overview of the FiberNeat method. Panel A shows whole-brain centroids and one expanded cluster of streamlines in DIPY’s viewer (A.a), an input cluster of streamlines (A.b), and visualization of their MDF distance matrix (A.c). Each streamline is mapped to a single 2D point using t-SNE (B.a) and clustered over the t-SNE embedding (B.b). Outlier streamlines are filtered out using our proposed approach (B.c).

FiberNeat requires two parameters, perplexity for t-SNE or n neighbors for UMAP, and epsilon for DBSCAN. For FiberNeat t-SNE, a smaller perplexity value gives attention to local structure(s) within a bundle, whereas a higher value of perplexity tries to preserve the global structure of bundles. It is non-trivial to find one value of perplexity that will work for different types of bundles with different shapes, sizes, and lengths. We empirically found an approach to automatically find perplexity *p*, and epsilon *eps* values depending on the number of streamlines *n* in the bundle. For small bundles with *n <*800, *p*=0.25 * *n* and *eps*=0.015 * *p*, for medium density bundles (most bundles belong in this category) with 800*< n <*4000, *p*=*n* * 0.065 and *eps*=*p* * 0.006, and for larger bundles such as the corpus callosum with *n >*4000, *p*=0.02 * *n* and *eps*=0.009 * *p*. For UMAP, n neighbors *p* is set to 0.05 * *n* for all types of bundles. *eps*=0.0025 * *p* and if *n <*800, *eps*=1.3. For most bundles, it takes less than 30 seconds to run FiberNeat. In Sec. III, we report individual timings, based on the type of bundle and the number of streamlines in a bundle.

## III. Results

In Fig.2, we show results on data from a 26-30 year-old male participant in the HCP (Human Connectome Project) [15], scanned with 90 diffusion weighting directions and 6 b=0 acquisitions. Diffusion weighting consisted of 3 shells of b=1000, 2000, and 3000 s/mm^2^. The tractogram was generated using deterministic local tracking. In Fig.2A, we show results on four clusters selected from all clusters in the whole-brain tractogram given by QuickBundles [6] with clustering threshold set to 25 mm. The first row shows the initial four clusters. The second row shows clusters cleaned manually by a trained neuroanatomist, using visualization tools in DSI Studio [16] and DIPY [17], [18]. We keep them as a ground truth to compare performance of FiberNeat t-SNE and FiberNeat UMAP. The third and fourth rows show clusters filtered using FiberNeat with t-SNE and UMAP embedding, respectively. In Fig.2B, a quantitative comparison of FiberNeat t-SNE and FiberNeat UMAP’s filtered clusters with expert cluster cleaning is shown. Here, SM stands for shape similarity score [9] among two clusters and BMD stands for bundle-based minimum distance [19] between clusters. SM scores range from 0 to 1, where 0 implies least shape similarity between two clusters/tracts and 1 means highest shape similarity. BMD calculates streamline-based distance between clusters in mm. A lower value of BMD implies that two clusters are closer and more similar in shape and streamline count. FiberNeat t-SNE’s filtered clusters have higher shape similarity and lower BMD distance relative to the expert’s cleaned clusters, except for cluster C3. FiberNeat UMAP’s output for C3 has higher shape similarity and lower BMD distance to an expert’s cleaned cluster C3. Overall, qualitatively and quantitatively, FiberNeat t-SNE performs better than FiberNeat UMAP.

**Fig. 2.**
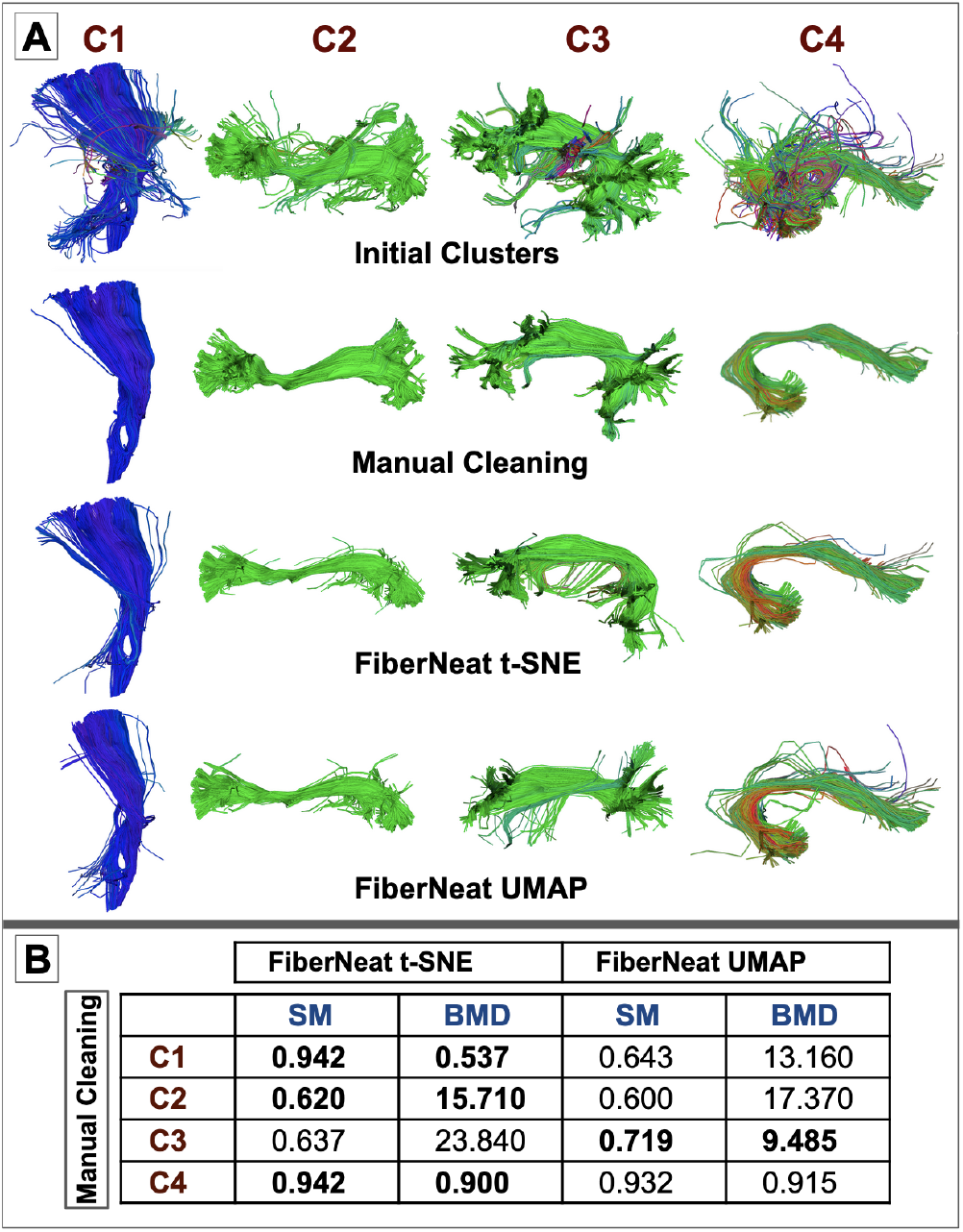
Part A, first row shows 4 initial clusters, the second row shows clusters manually cleaned by an expert. The third and fourth rows show clusters cleaned by FiberNeat t-SNE and FiberNeat UMAP, respectively. Part B shows the quantitative comparison of FiberNeat t-SNE and FiberNeat UMAP clusters with expert’s cleaned clusters. Shape similarity score (SM) and bundle minimum distance (BMD) are calculated between clusters.

To test the algorithms’ performance on a larger dataset, we analyzed whole-brain tractography computed from multi-shell diffusion MRI (dMRI) data from 131 Alzheimer’s Disease Neuroimaging Initiative phase 3 (ADNI3) [20] participants (age: 55-91 years, 74F, 57M) scanned on 3T Siemens scanners. dMRI consisted of 127 volumes: 13 b0, 48 b=1,000, 6 b=500 and 60 b=2,000 s/mm^2^ volumes with an isometric 2-mm voxel size. Participants included 44 with mild cognitive impairment (MCI) and 87 cognitively normal controls (CN). dMRI were preprocessed using the ADNI3 dMRI protocol, correcting for artifacts including noise, Gibbs ringing, eddy currents, bias field inhomogeneity, and echo-planar imaging distortions [21]. We applied multi-shell multi-tissue constrained spherical deconvolution [22] and a probabilistic particle filtering tracking algorithm [23] to generate whole-brain tractograms. We extracted 30 white matter tracts from tractograms using auto-calibrated RecoBundles [8], [9]. In Fig.3, we show a use case of FiberNeat as a spurious streamline filtering method deployed after the bundle extraction method. In this experiment, we used RecoBundles (RB) [8] to extract white matter tracts and used FiberNeat on its output to eliminate any spurious streamlines. RB takes a model bundle as a reference and tries to extract similar looking streamlines from the input tractogram. We visually illustrate the results on one of the ADNI3 subjects. The first row shows model bundles for four tracts: the arcuate fasciculus (AF L), middle longitudinal fasciculus (MdLF L), Uncinate Fasciculus (UF L), and Optic Radiation (OR L) in the left hemisphere of the brain. The second row shows the output bundles from RB. RB output bundles were given as input to FiberNeat. The third row shows the output of FiberNeat. The fourth row visualizes the overlap between RB output and FiberNeat output. Red streamlines are the ones filtered out by FiberNeat from the RB bundles. As compared to the experiment in Fig.2, Fig.3’s experiment starts with cleaner input tracts and only removes outlier streamlines as seen in AF L, and UF L tracts. FiberNeat does not remove streamlines unnecessarily, as can be seen in the case of OR L. It only removes a few spurious streamlines from OR L. From the MdLF L bundle, it removes a cluster that is crossing the actual bundle fibers and might have been mistakenly labeled as part of the bundle by the tract extraction method. FiberNeat MdLF L matches the model bundle.

**Fig. 3.**
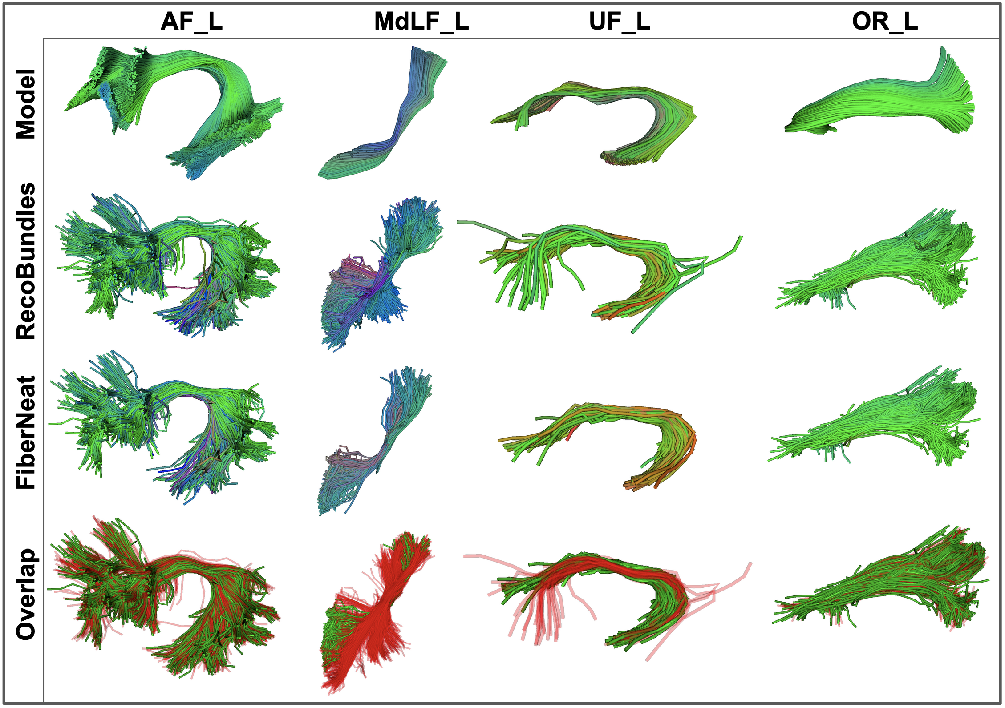
Four columns representing four different bundles. The first row shows a model bundle used in RecoBundles. The second row shows the RecoBundles output. The third row shows the FiberNeat output, with the RecoBundles output used as input. The fourth row shows outliers in red, that were removed by FiberNeat from RecoBundles’ output.

FiberNeat is a fast method and takes around 30 seconds to run for most bundles. Since bundles have different lengths and sizes, the time it takes to run FiberNeat depends on the number of streamlines in the bundle. Referring to bundles in Fig.3, for smaller bundles such as UF L with 220 streamlines, it took 1.10 seconds to run FiberNeat. Most bundles have streamlines in the range of 800-4000 - such as OR L with 987, AF L with 1331, and MdLF L with 1934 streamlines. On these, it took 11.36, 18.00, and 37.55 seconds respectively to run FiberNeat. However, for larger bundles such as Corpus Callosum (CC) with more than 4000 streamlines, runtime is longer as it has to calculate distances among all the streamlines.

We performed bundle shape similarity analysis on tracts extracted from 131 subjects. We calculated shape similarity among 30 model bundles and 30 RB extracted tracts per subject and separately among 30 model bundles and 30 FiberNeat cleaned RB tracts. In Fig.4, we visualize results from two experiments. A) and B) plots both show a 30×131 shape similarity plot where rows have a shape similarity score between model bundles used as a reference in RB and extracted bundles of the same type by RB (A) and FiberNeat cleaned RB bundles (B). Columns of the plots represent the 131 subjects. Each pixel is a bundle shape similarity score between the model bundle and extracted bundle from a subject. B) plot is darker than the top plot indicating higher shape similarity among model bundles and FiberNeat bundles. In some subjects and bundles, we see less shape similarity after FiberNeat. This could be due to FiberNeat cleaning and making some bundles very thin. Overall, we observe that bundle shape similarity tends to improve after deploying the FiberNeat step after RB. We ran paired t-test on shape similarity scores from 131 subjects per bundle to test the hypothesis that FiberNeat improves shape similarity among bundles. We ran false discovery rate (FDR) on p-values and report results in C). In the top plot in C), we observe significant improvement in bundle shape similarity after using FiberNeat for all bundles except the frontopontine tract in both left and right hemispheres (FPT L, and FPT R). The bottom plot in C) shows average RB and FiberNeat shape similarity scores per bundle. On the right end of the figure, 30 bundle names are listed.

**Fig. 4.**
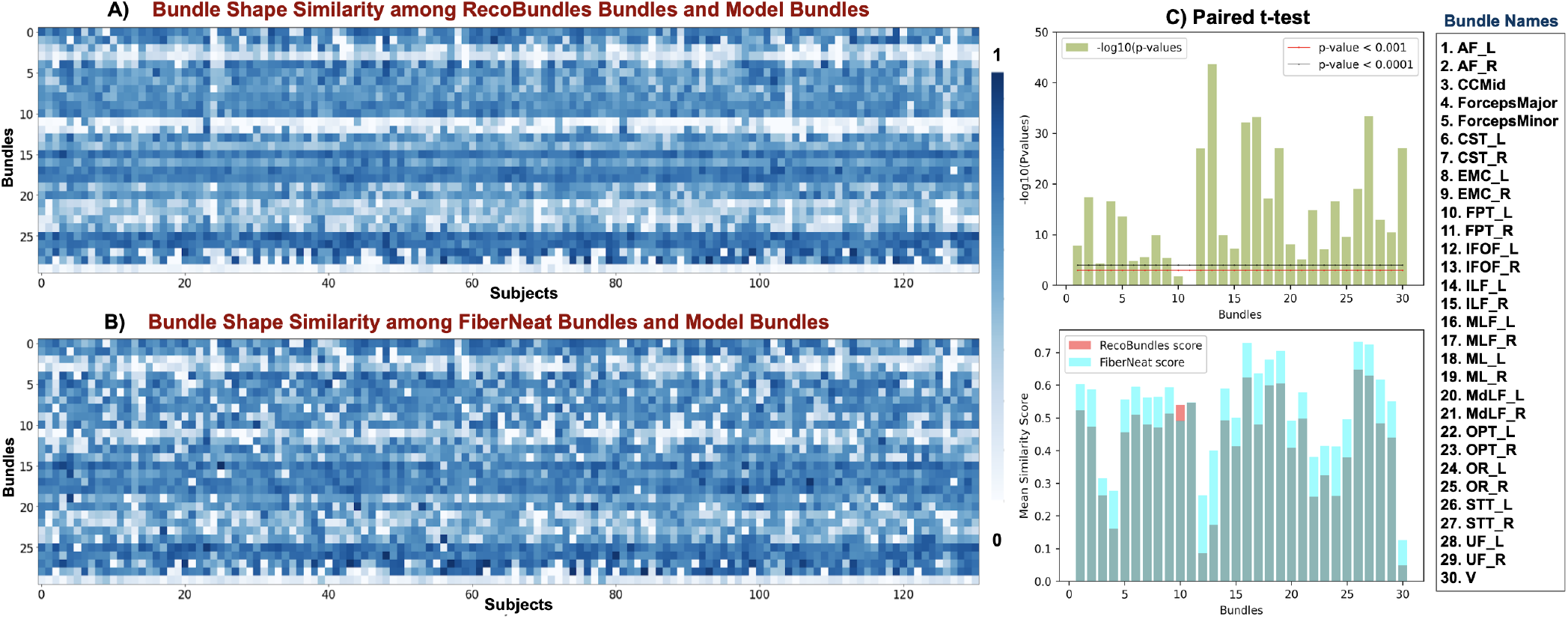
A) Bundle shape similarity among RecoBundles bundles and model bundles. B) Shape similarity among FiberNeat bundles and model bundles. In both 30×131 matrices, the x-axis has subject numbers and the y-axis has bundle numbers labeled on it. Each pixel is a bundle shape similarity score between model bundle and RecoBundles output bundle (A) and score between model bundle and FiberNeat output bundle (B). C) Negative logarithm of p-values from a paired t-test to test FiberNeat improves shape similarity scores (top), mean bundle shape similarity score per bundle from two experiments (bottom). On the right panel, we provide the names of the 30 bundles used here.

We ran along the length tract group analysis of fractional anisotropy (FA), a commonly used white matter microstructural measure, between groups of 44 MCI and 87 CN participants using BUndle ANalytics (BUAN) [9]. We ran BUAN twice, once with RB bundles and the FA metric as input and another time by applying FiberNeat on RB bundles and using filtered bundles and the same FA metric as input to BUAN. In Fig.5, we show results for the MdLF L bundle. BUAN creates 100 horizontal segments along the length of the bundles and analyzes points on the streamlines belonging to each segment from all subjects. It then applies Linear Mixed Models (LMMs) where group means are modeled as a fixed effects term and the subject-specific mean is modeled as a random effects term with FA as a response of the LMMs. BUAN plots in Fig.5 have segment number on the x-axis and a negative logarithm of p-values on the y-axis. P-values that lie between or above two horizontal lines on the plot imply significant group differences at that location along the tract. We find that by deploying FiberNeat on the output of RB, we are able to improve the tractometry by removing spurious streamlines that can cause artifacts in the group analysis. Areas on the bundles that FiberNeat cleaned are indicated by arrows in the Fig.5. Red arrows indicate segments between 20-40 on the bundles. Those areas show significant p-values when BUAN is run on RB output and significance reduces slightly after removing outlier streamlines using FiberNeat (last plot). Blue arrows indicate segments around 80-100. By removing outliers from that area on the bundle, we observe stronger significant group differences in FA (lower p-values). By removing streamlines of different shapes, we focus on the same type of streamlines and are able to better find the effects of MCI on the MdLF L tract.

**Fig. 5.**
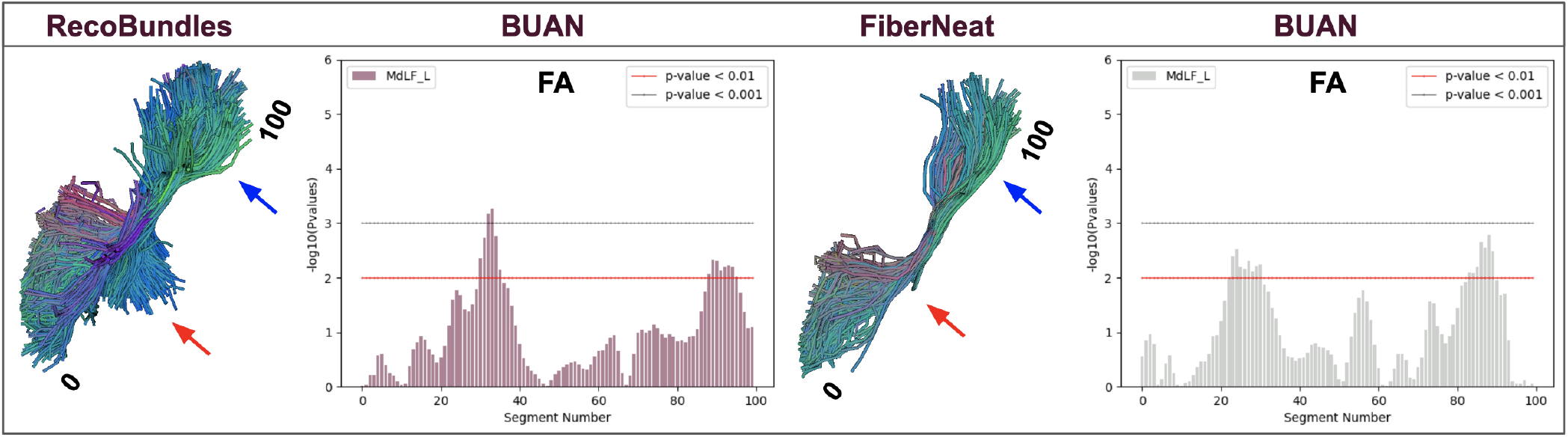
Improved tractometry with FiberNeat. On the left, BUAN tractometry results on MdLF L tract using RecoBundles output. On the right, BUAN tractometry results on RecoBundles’ MdLF L tracts cleaned by FiberNeat. FiberNeat removes spurious streamlines that might cause artifacts in the group analysis of FA microstructural differences along the length of the tracts in MCI and CN groups (indicated by arrows).

## IV. Discussion

Tractography data is unstructured complex data, and sub-dividing it into bundles is a highly nonlinear problem. It is also difficult to perfectly separate outliers from good clusters corresponding to known anatomical tracts. t-SNE and UMAP are both manifold learning approaches for non-linear dimensionality reduction. Clustering based on t-SNE and UMAP embedding of tractography makes it easier to separate streamline clusters and outliers. Some researchers caution against clustering the t-SNE embedding space, at least for some applications, due to metric distortions. t-SNE is a stochastic method and can generate different embeddings in different runs for the same data and parameters. It does not preserve the global metric structure and favors the preservation of the local structure only. t-SNE can sometimes disconnect/split parts of the data by putting them in separate clusters. This repelling effect of t-SNE is advantageous in our application as we want to untangle streamlines that are otherwise very closely knitted together in the original space, as seen in Fig.1C.a. The stochastic nature of t-SNE does not affect our approach as we do not use the embedding map again, for further data analytics. It is used once per input dataset and the method is invariant to where clusters are placed and to the global distance among clusters. t-SNE does extreme dimensionality reduction by going directly to 2D space as opposed to other dimensionality reduction methods that provide options to project data into n*>*2 dimensions. But in our case, every streamline has *k* points and each point is a 3D vector making it *k* × 3 D, and going to 2D is not an extreme dimensionality reduction. We also provide an option to use UMAP embedding instead of t-SNE. Theoretically, UMAP should give superior performance relative to t-SNE. UMAP tries to preserve both local and most of the global structure in the data. UMAP can map data to latent spaces with any number of dimensions and does not need the pre-dimensionality reduction step such as PCA or an autoencoder. Hence, UMAP can project data on n components and is not limited to 3D or 2D embeddings (as required by t-SNE). UMAP is computationally faster than t-SNE. However, in our experiments, we find t-SNE to outperform UMAP. This could be because the nature of the problem we are solving takes advantage of the data splitting/repelling property of t-SNE to find outlier streamlines in streamline sets that hard to distinguish in the original brain’s 3D space. Further work is needed to evaluate the method on more tracts, and diverse datasets (in terms of age, diagnosis, and scanning protocol). We keep t-SNE as the default dimenrsionality reduction method in FiberNeat, however, users can select UMAP as well. The main advantage of the Fiberneat approach is that it is fully data-driven and unsupervised. It does not depend on training data or an atlas as compared to recently proposed methods for tractography filtering using deep learning [24], [25].

We show two use cases of FiberNeat, one applied to the output of unsupervised clustering-based bundle segmentation method, QuickBundles (QB), and the other applied to the output of supervised bundle segmentation method, RecoBundles (RB). For QB, the clustering threshold set was 25 mm. Decreasing the threshold may result in more cluters with less spurious streamlines in them. However, in out experiments, we show even with densely populated clusters, FiberNeat is able to perform well, comparing results with expert delineated clusters. For RB, we used an auto-calibrated version with a pruning threshold of 12 mm and a refine-pruning threshold of 9 mm. We used moderate threshold values for auto-calibrated RB in our experiments, reducing thresholds could result in thin bundles and less spurious streamlines but we might also risk losing some parts of bundles. RB uses prior anatomical information from the model bundle, whereas FiberNeat only uses the input bundle to clean it, without using any external or prior information. RB performs well extracting bundles, especially for larger or medium sized bundles. It sometimes extracts spurious streamlines for smaller bundles with average bundle length of 50 mm such as the uncinate fasciculus. This could happen due to some streamlines having very small length in those bundles and it becomes difficult to distinguish which streamlines match and which do not match with a model bundle as the distance between streamlines is small, even if they have different shapes. Diffusion MRI and tractography provide crucial information about the brain connectivity and microstructural changes in it due to any underlying condition such as Alzheimer’s, Parkinson’s, Schizophrenia, etc. Tractometry tools to study microstructural changes along the length of white matter tracts have been gaining popularity in the past decade. Several tractometry methods have been proposed and used to study brain diseases and to find group differences in patients and healthy controls [26], [27], [28], [9]. These methods rely on bundle segmentation methods and false-positive streamlines extracted in the output bundles could propagate artifacts in statistical analysis of microstructural measures along the length of the tracts. In this paper, we used BUAN to find group differences in FA measure along the length of MdLF L tracts of 87 CNs and 44 participants with MCI from the ADNI3 dataset. We find that having spurious streamlines in input data can overestimate or underestimate the effects of the disease. We show deploying FiberNeat into BUAN tractometry pipeline improves the robustness of statistical analysis by removing spurious and false-positive streamlines that could create artifacts in the analysis. We show adding FiberNeat as a cleaning step into BUAN tractometry pipeline improves the robustness of statistical analysis by removing any artifacts introduced by outlier streamlines as shown in Fig.5. FiberNeat code and tutorial are available here: https://github.com/BramshQamar/FiberNeat. It will also be made available through DIPY [17].

## V. Conclusion

In this paper, we introduce FiberNeat, a method to clean streamline clusters and tracts. It takes a set of streamlines as input, calculates a distance matrix of all the pairwise streamline distances, and projects it into a reduced low-dimensional space using dimensionality reduction techniques such as t-SNE and UMAP. FiberNeat applies DBSCAN clustering in 2D space. Smaller clusters containing spurious streamlines are removed from the original data in the streamline space resulting in cleaner tracts. We tried FiberNeat with t-SNE and UMAP on several same clusters of streamlines and found FiberNeat t-SNE to perform better than FiberNeat UMAP, both qualitatively and quantitatively. We used FiberNeat on the output of the unsupervised clustering method, QuickBundles, and on the output of the supervised bundle segmentation method, RecoBundles. In both experiments, FiberNeat removes spurious and outlier streamlines and improves the quality of final clusters/tracts. We propose FiberNeat to be used as a post-processing step in tract segmentation methods to aid better tractometry. We show on 131 ADNI3 participant data (Fig.5) that deploying FiberNeat after tract segmentation reduces false-positive streamline artifacts that could propagate in statistical analysis of microstructural measures along the length of the white matter tracts.

## ACKNOWLEDGMENT

This research was supported by the NIH (National Institutes of Health) under the AI4AD project grant U01 AG068057, grant numbers P41 EB015922, and RF1 AG057892. We would like to acknowledge the National Institute of Biomedical Imaging and Bioengineering under award number R01 EB027585. Paul M Thompson received a research grant from Biogen, Inc. for research unrelated to this manuscript.

## Footnotes

1 Mapping high-dimensional data to a latent space refers to transforming complex forms of raw data into a simpler, lower-dimensional representation

